# Structural studies of the C-terminal cysteine-rich domain of *Aedes albopictus* vitellogenin reveal an evolutionarily divergent, monomeric C-terminal cysteine knot (CTCK) architecture

**DOI:** 10.64898/2026.05.25.727680

**Authors:** Marco Berlinguer, Mahdieh Sadeghichehelgaz, Alice Vetrano, Alessandro D’Aquilio, Ferdinando Mercuri, Marta Villa, Paolo Gabrieli, Claudio Iacobucci, Federico Forneris

## Abstract

Vitellogenins are essential transport lipoproteins and precursors to egg-yolk proteins in oviparous species. Several molecular structure studies have elucidated most of their multi-domain organization, yet the structure and function of their C-terminal cysteine knot (CTCK) domain have remained largely elusive. In this study, we present the 1.2 Å resolution crystal structure of the recombinant CTCK domain from a vitellogenin isoform of the mosquito *Aedes albopictus* (Vg-CTCK). The molecular architecture reveals a CTCK fold defined by two antiparallel beta sheets stabilized by three intramolecular disulfide bonds, featuring a unique 12-amino acid insertion shaping an alpha helix positioned between the two main beta sheets.

Analysis of the crystal packing and biophysical characterization in solution consistently confirms that recombinant Vg-CTCK is monomeric. To validate these findings in a native context, we employed *ex vivo* cross-linking mass spectrometry (XL-MS) on intact mosquito ovaries, which corroborated the molecular architecture of the Vg-CTCK observed in the crystal structure and highlighted the absence of inter-molecular cross-links.

Collectively, our data highlight an evolutionarily divergent, monomeric assembly for the mosquito Vg-CTCK domain, challenging previous hypotheses that suggested this domain might facilitate vitellogenin oligomerization.

## Introduction

Vitellogenins are large transport glycolipoproteins present in most oviparous living beings [1]. These macromolecules play a fundamental role in egg maturation, as they constitute the major precursors of egg-yolk proteins. In vertebrates, vitellogenins are produced in the liver, whereas in insects and invertebrates their synthesis occurs in a functionally analog organ, such as the fat body and or the follicular cells in the ovaries.

In honeybees, vitellogenin synthesis is regulated by Juvenile hormone levels and shows differences between workers and queen’s castes [2,3]. In contrast, mosquito vitellogenin genes in the female fat body he concerted action of nutritional and hormonal stimuli, specifically the TOR signaling pathway, the insulin-like peptides, the juvenile and the 20-hydroxyectdisone (20E) hormones [4] and therefore, by the blood feeding behavior. Few hours after the blood meal, female mosquitoes undergo a vitellogenesis peak right before egg maturation and oviposition [5-9]. The fat body releases mature vitellogenins into hemolymph, which pass to oocytes diffusing through channels between cells forming the follicular epithelium. They are internalized by a process of receptor-mediated endocytosis, binding vitellogenin receptors located in coated pits on the oolemma [10].

Molecular studies of vitellogenins have leveraged on early X-ray crystallography characterizations of lipovitellins and phosvitin assembly (i.e., post-internalization proteolytically-processed vitellogenins) from silver lamprey [11]. Recently, the molecular architecture of a honeybee vitellogenin prior to internalization was determined, using cryo-electron microscopy on purified haemolymph from wintering bees [12]. In all cases, the molecular structures were determined at near-atomic resolution, enabling accurate mapping of the amino acid networks composing most domains shaping the complex vitellogenin three-dimensional architecture. Indeed, vitellogenin polypeptides are characterized by a multi-domain architecture encompassing a large N-terminal beta barrel domain, responsible for interactions with the vitellogenin receptor [13], followed by a alpha helix solenoid domain that wraps around multiple beta sheet domains. The C-terminus is characterized by a Von Willebrand factor type D domain and a C-terminal cysteine knot domain (CTCK), which has remained particularly elusive until now.

Proteins belonging to the CTCK domain family are typically peptides shorter than 100 amino acids who presents two or three intramolecular disulfide bonds that stabilize the folding. CTCK domains are typically found in extracellular growth factors and modular proteins such as the Von Willebrand factor (VWF), and are characterized by a well-defined cluster of disulfide bonds interlinking the N- and C-terminal segments of the domains, often resulting in inter-molecular linkage contributing to protein dimerization [14,15]. Several hypotheses regarding the function of Vg-CTCK have been made [12,16], mostly based on theoretical modeling and sequence similarity, but without experimental support. In this context we report the recombinant expression, characterization, and high-resolution structure determination of the CTCK domain of a Vitellogenin isoform from *Aedes albopictus* (Vg-CTCK).

## Materials and Methods

### Molecular Cloning

A codon-optimized cDNA construct encoding *Aedes albopictus* Vitellogenin gene (NCBI entry XP_019542945.3, corresponding to UniProt entry A0ABM1Z0L2), corresponding to the C-terminal domain, from residue 1998 to end, was synthetized (GeneWiz) and cloned into the a modified pET-SUMO recombinant expression plasmid (Life Technologies) using in frame 5’-BamHI and 3’-NotI restriction sites. The resulting protein construct encodes for 8xHis–SUMO-Vg-CTCK.

### Recombinant protein production

The recombinant pET-SUMO-Vg-CTCK plasmid was transformed into *Escherichia coli* T7 SHufsle chemically competent cells (New England Biolabs). A single colony was used to inoculate 200 mL of Lysogeny medium, supplemented with 0.1 mg/mL kanamycin. This pre-culture was incubated overnight at 30 °C in a shaking incubator at 180 rpm. The following day the pre-culture was used to inoculate 2 liters of autoinducing ZYP-5052 medium [17] supplemented with 0,1 mg/mL kanamycin, for large scale protein expression. Cultures were grown for 4 h at 30 °C, then the temperature was decreased to 18 °C and incubation continued for additional 20 h prior to cell harvesting. The bacterial cells were harvested by centrifugation at 5000 g, 10 °C for 20 minutes. The bacterial cell pellet was resuspended in buffer A (25 mM HEPES/NaOH, 500 mM NaCl, pH 8) using a 1:10 (*w/v*) wet cell pellet:buffer ratio. Cells were then disrupted by sonication (20 cycles, 9 seconds on, 9 seconds off pulses). Cell debris were removed by centrifugation at 40’000 g, 10 °C for 45 minutes. The supernatant was filtered through a 0.8 μm syringe-driven filter (MiniSART GF, Sartorius) and loaded onto a 5 mL prepacked His-Trap Excel column (Cytiva), pre equilibrated with buffer A at a flow rate of 1 mL/min on a Åkta Go chromatography system (Cytiva). The column was first washed with buffer A supplemented with 25 mM Imidazole for a total of 10 CV to remove non-specific binding contaminants. Then the elution of target protein was performed by using buffer A supplemented with 250 mM imidazole. The elution fractions containing the protein of interest, assessed by SDS-PAGE, were pooled together, supplemented with SUMO protease and dialyzed overnight at 4°C. After dialysis the sample was centrifuged (15 minutes, 5000 g, 10°C) and reloaded onto the 5 mL prepacked His-Trap excel column (Cytiva) pre equilibrated with buffer A at a flow rate of 1 mL/min. Cleaved Vg-CTCK eluted in the flow-through fraction. The sample was concentrated using a 3 kDa Amicon Ultra-15 centrifugal filter (Merck) and further purified via gel filtration using a Superdex 75 10/300 GL Increase column (Cytiva) pre-equilibrated with SEC buffer (25 mM HEPES/NaOH, 200 mM NaCl, pH 7,5). Vg-CTCK containing fractions were pooled together. Sample purity was assessed by SDS-PAGE before flash freezing in liquid nitrogen for further usage.

### Crystallization, structure determination and refinement

Purified Vg-CTCK was thawed, concentrated to 22 mg/mL and subsequently screened for crystallization against the sparse-matrix commercial screening kit JCSG+ (Molecular Dimensions). Crystallization drops were prepared using a Gryphon crystallization robot (Art Robbins Instruments) and kept at 4 °C. After few, days, crystals deemed suitable for diffraction experiments were found in conditions containing 0.1 M HEPES/NaOH, 30% v/v Jeffamine ED-2003, pH 7.0. Cryoprotection was achieved by adding 20% (v/v) glycerol to the crystallization mother liquor. Crystals were harvested using mounted Litholoops (Molecular Dimensions), flash-cooled in liquid nitrogen, and sent to the European synchrotron radiation facility (ESRF). X-ray diffraction data were collected at beamline ID30A-3. Data were indexed and integrated with *XDS* [18], followed by scaling and merging using *AIMLESS* [19]. Data collection statistics are given in Table 1. Structure determination was carried out by molecular replacement with *Phaser* [20], using a computational model of Vg-CTCK generated using Alphafold3 [21] as search model. Structure refinement was carried out by alternating steps of manual building in *COOT* [22] and automated refinement with *phenix*.*refine* [23] and *REFMAC5* [24]. Validation was carried out using *MolProbity [25]* and the validation tools available on the Protein Data Bank server [26]. Final refinement statistics are summarized in Table 1. Structural figures were prepared with *PyMol* [27] and *ChimeraX* [28]. Superpositions were performed using the “*super*” or the “*cealign*” commands in *PyMol*. All-atom root mean square deviation (RMSD) values were computed accordingly.

**Table 1.**
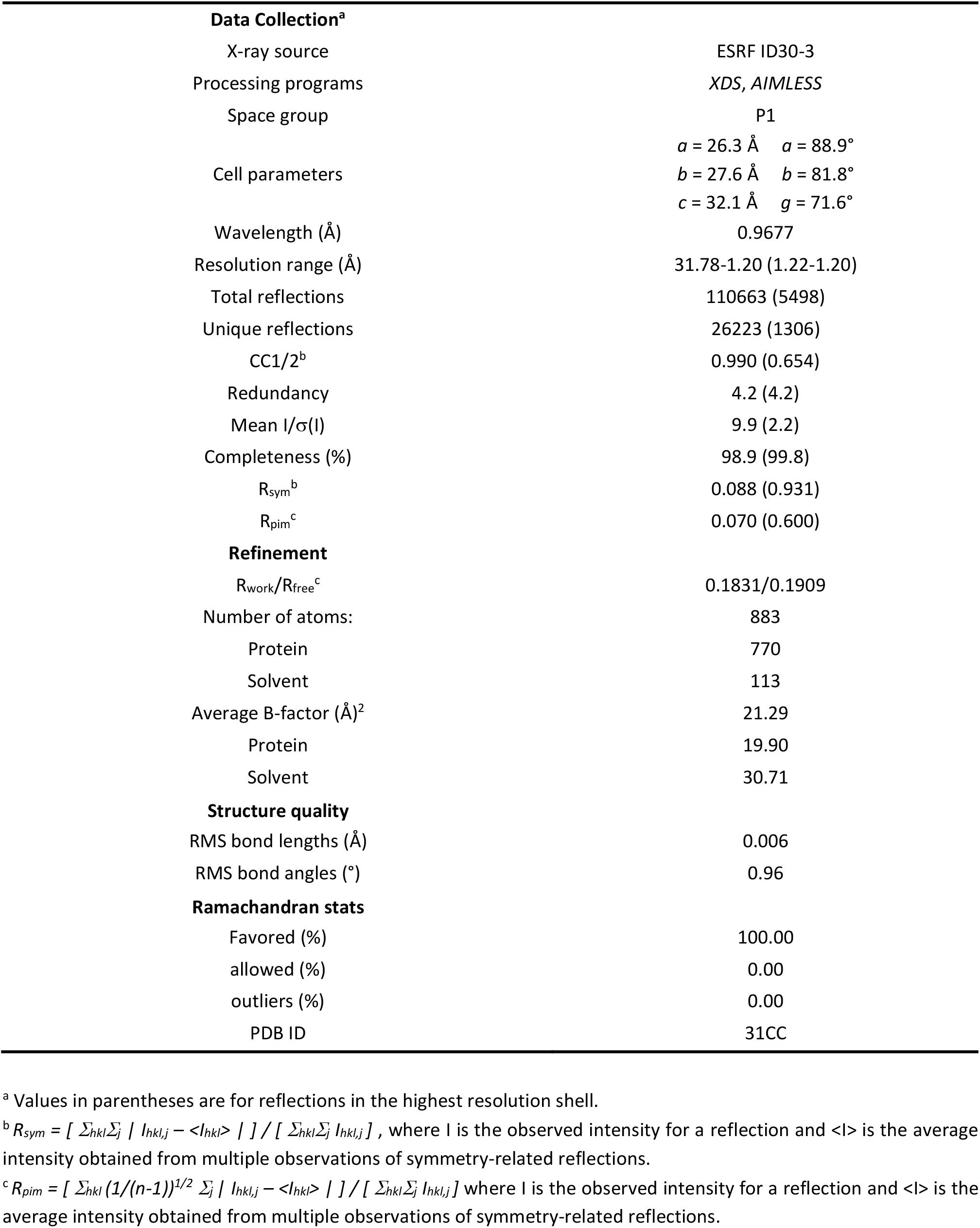
X-ray diffraction data collection and refinement statistics.

### Size exclusion chromatography coupled to multi-angle light scattering (SEC-MALS)

40 ml of 5 mg/mL recombinant Vg-CTCK were injected into a Protein KW-802.5 analytical size-exclusion column (Shodex) and separated with a flow rate of 1 mL/min in phosphate buffer saline using a Prominence high-pressure liquid chromatography (HPLC) system (Shimadzu). For molecular weight characterization, light scattering was measured with a miniDAWN multi-angle light scattering detector (Wyatt), connected to a RID-20A differential refractive index detector (Shimadzu) for quantitation of the total mass and to a SPD-20A UV detector (Shimadzu) for evaluation of the sole protein content. Chromatograms were collected and analyzed using the *ASTRA7* software (Wyatt), using an estimated *dn*/*dc* value of 0.185 ml/g). The calibration of the instrument was verified by injection of 10 mL of 2.5 mg mL^-1^ monomeric BSA.

### Ex vivo ovaries cross-linking

*Aedes albopictus* females were collected from the breeding colony at the Department of Biosciences (Università degli Studi di Milano, Italy). This strain was maintained under standard insectary conditions (27 ± 1 °C, 65–80% relative humidity, 12:12 h light/dark photoperiod). Larvae were fed daily with carp fish pellets (Tetramin, Tetra), pupae were separated in a small cup of water and moved to adult cages (Bugdorm). Adults were maintained on a 10% sucrose solution. About 30 mature females (6-7 days) were fed on human volunteer arm (Ethical Committee of the University of Milan 13.22). After 24 hours, when females reach the peak of vitellogenesis, the ovaries of 15 females were dissected in Schneider’s medium and collected by centrifugation at 10,000 × g for 3 min. The pellet was resuspended in PBS containing either 0.5 or 1 mM DSBU and incubated for 1 h at 25 °C to allow cross-linking. Excess DSBU was quenched by addition of Tris-HCl, pH 8.0, to a final concentration of 10 mM. Samples were then snap-frozen in liquid nitrogen.

### Sample Processing and Digestion

Cross-linked ovary samples were centrifuged and resuspended in 46 μL of 50 mM Tris buffer (pH 8.5) containing 5% sodium dodecyl sulfate (SDS) and sonicated until complete solubilization. Disulfide bonds were reduced with 5 mM TCEP (Merck) at 55 °C for 15 min and subsequently alkylated with 20 mM iodoacetamide (Merck) in the dark for 10 min. The solution was then acidified by adding phosphoric acid to a final concentration of 1.2%. Proteins were precipitated as fine particles by the addition of 90% methanol in 100 mM Tris buffer (pH 7.55). The resulting suspension was loaded onto an S-Trap Micro column (Protifi) and centrifuged at 4,000 × g until the entire volume passed through the protein-trapping matrix. The captured proteins were washed three times with 150 μL of 100 mM Tris buffer (pH 7.55) in 90% methanol. Digestion was initiated by adding 20 μL of 50 mM Tris buffer containing Trypsin Gold (Promega) at an enzyme-to-protein ratio of 1:25. Proteolysis was performed overnight at 37 °C, followed by a 2 h incubation at 47 °C. Resulting peptides were sequentially eluted using: (i) 50 mM Tris buffer (pH 8.5), (ii) 0.2% formic acid in water, and (iii) 50% acetonitrile in water. The eluates were pooled and vacuum-dried.

### SEC fractionation

200 ug of peptides obtained after tryptic digestion of cross-linked proteins were dried under vacuum and resuspended in size exclusion chromatography (SEC) buffer consisting of 30% (v/v) acetonitrile and 0.1% (v/v) formic acid in water. Prior to injection, samples were briefly vortexed and centrifuged to remove insoluble material. SEC fractionation was performed using a Superdex 30 Increase 3.2/300 (Cytiva) column coupled to an Agilent 1100 Series high-performance liquid chromatography (HPLC) system equipped with a diode array detector (DAD). The separation was carried out under isocratic conditions using the SEC buffer as mobile phase at a flow rate of 0.05 mL/min, for a total run time of 60 min. Elution was monitored by UV absorbance at 280 nm, and fractions were collected every 1 min, resulting in 30 fractions per run. Since cross-linked peptides are expected to have a larger hydrodynamic radius than linear peptides, six 50 µL early-eluting fractions (between 32 and 37 min) were selected and dried for downstream LC–MS/MS analysis.

### Mass Spectrometry. Nano-HPLC/Nano-ESI-MS/MS Analysis of Cross-Linked Samples

Peptides before and after SEC fractionation were resuspended in 0.1% (v/v) trifluoroacetic acid (TFA) in water and analyzed by nanoLC-MS/MS using a Vanquish Neo UHPLC system (Thermo Scientific) coupled to an Orbitrap Eclipse Tribrid mass spectrometer (Thermo Scientific) equipped with a Nanospray Flex ion source (Thermo Scientific) and a 30 μm ID steel emitter (Thermo Scientific). Peptides were separated on a PepMap Neo column (75 μm × 500 mm, 2 μm particle size, 100 Å pore size; Thermo Scientific, San Jose, CA). The gradient consisted of 2% phase B (80% acetonitrile with 0.1% formic acid) for 15 min, followed by a linear increase to 37% B over 240 min, then ramped to 50% and 99% B in two consecutive 5 min steps, followed by column washing and re-equilibration. Data were acquired in data-dependent acquisition (DDA) mode with a 3 s cycle time. Full MS scans were collected over an m/z range of 450–1300 in the Orbitrap (resolution: 120,000 at m/z 200) with a maximum injection time of 50 ms and an AGC target of 8 × 10^5^. Dynamic exclusion was set to 30 s, selecting precursor ions with charge states from 2+ to 6+. Peptides were fragmented using stepped HCD at 26%, 29%, and 32% normalized collision energy, and MS/MS spectra were acquired at a resolution of 60,000.

### Cross-Link Data Analysis

Raw LC–MS/MS data were processed using Proteome Discoverer (version 3.1). Spectra were exported in Mascot Generic Format (MGF) files. Protein identification was performed using the *SEQUEST HT* search engine with Percolator against the *Aedes albopictus* genome (AalbF5) protein database (downloaded from NCBI, release January 4, 2024), supplemented with common contaminants. Search parameters included a precursor mass tolerance of 10 ppm and a fragment ion mass tolerance of 0.02 Da. Enzyme specificity was set to trypsin, allowing up to 3 missed cleavages. Carbamidomethylation of cysteine residues was set as a fixed modification, while oxidation of methionine, acetylation of the protein N-terminus, and DSBU hydrolysis were considered as variable modifications. Based on protein identification results, the 30 most abundant proteins were selected for subsequent cross-linking analysis, including three vitellogenins. Identification of cross-links was performed using *MeroX* 2.0.2.5 [29] and Scout 2.1.0 [30]. The following settings were applied in *MeroX*: proteolytic cleavage C-terminal to Lys and Arg (up to three missed cleavages allowed); peptide length ranging from 5 to 30 amino acids; modifications included carbamidomethylation of Cys (fixed) and oxidation of Met (variable); cross-linker specificity was set to Lys, Ser, Thr, and Tyr; the RISEUP algorithm was used with a maximum of three missing ions; precursor mass tolerance was set to 10 ppm and fragment ion mass tolerance to 20 ppm; signal-to-noise ratio > 2; precursor mass correction was enabled; the false discovery rate (FDR) was controlled at 1% at the residue-pair level, with a minimum score cut-off of 20. The following settings were applied in Scout: proteolytic cleavage C-terminal to Lys and Arg (up to three missed cleavages allowed); peptide length ranging from 5 to 30 amino acids; modifications included carbamidomethylation of Cys (fixed) and oxidation of Met (variable); cross-linker specificity was set to Lys, Ser, Thr, and Tyr; precursor mass tolerance was set to 10 ppm and fragment ion mass tolerance to 20 ppm; the false discovery rate (FDR) was controlled at 1% at the residue-pair level. The results obtained from the two software tools were merged and visualized with *xiVIEW* [31].

## Results

### Recombinant production of Vg-CTCK from Aedes albopictus yields monomeric species in solution

*Aedes albopictus* (the Asian tiger mosquito) expresses multiple vitellogenin isoforms that display tissue-specific expression and regulation, in contrast to honeybees that possesses a single vitellogenin gene. In this work, we chose to focus on the isoform corresponding to NCBI entry XP_019542945.3 and UniProt entry A0ABM1Z0L2. A recombinant protein construct corresponding to the residues 1998-2077 of this entry, corresponding to the predicted structured portion of the C-terminal CTCK domain (Vg-CTCK), was generated. Based on sequence analysis, Vg-CTCK incorporates three of the canonical CTCK cysteine pairs, which are predicted to contribute to the stabilization of the native folding [15]. Vg-CTCK was successfully produced in a soluble form using an expression vector that encodes a 8xHis-SUMO fusion tag, which was then removed yielding the sole purified protein fragment in solution. Vg-CTCK eluted as a sharp symmetric peak in gel filtration well separated from other protein species, consistent with a stable, well folded globular protein fragment (Figure 1A). Sample purity was further assessed by SDS-PAGE in both reducing and non-reducing conditions, revealing a comparable migration pattern in both conditions (Figure 1B). SEC-MALS analyses carried out using recombinant Vg-CTCK consistently confirmed the presence of monomeric species in solution (Figure 1C).

**Figure 1.**
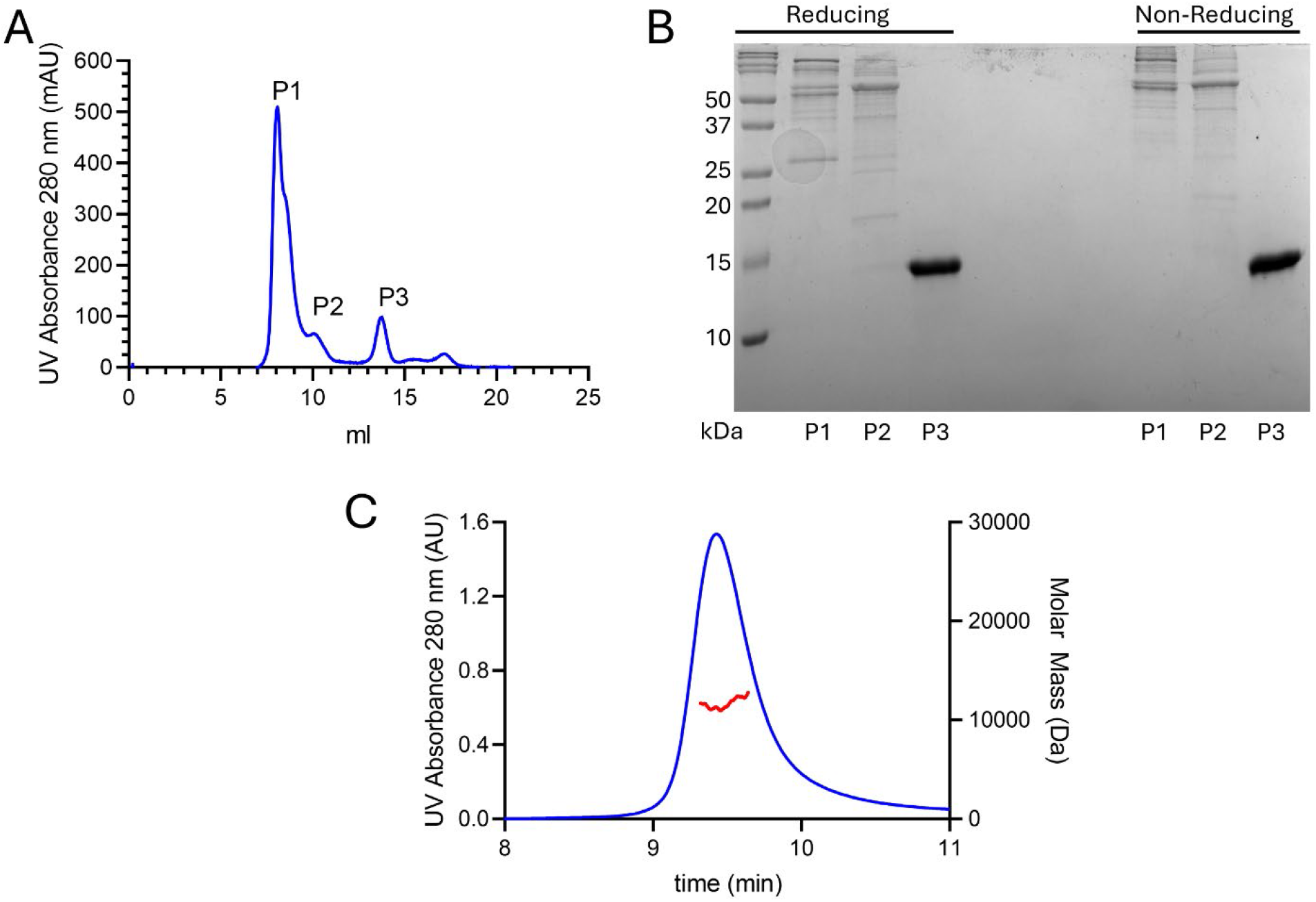
Recombinant production of *Aedes albopictus* Vg-CTCK. (A) Gel filtration chromatogram obtained at the end of recombinant Vg-CTCK purification. (B) SDS-PAGE analysis of purified Vg-CTCK samples in reducing and non-reducing conditions. The lanes include samples matching the contents of the peaks represented in the chromatogram shown in panel (A). (C) SEC-MALS analysis of Vg-CTCK.

### Crystal structure of Vg-CTCK

Crystals obtained in a reservoir containing 0.1 M HEPES/NaOH, 30% v/v Jeffamine ED-2003, pH 7.0 produced diffraction patterns extending to detector limit of beamline ID30A-3 at the ESRF, ultimately yielding a complete 1.2 Å resolution dataset. Structure determination was performed by molecular replacement, using a AlphaFold3-derived computational prediction as search model. The resulting phased electron density map was excellently defined for most of the residues of Vg-CTCK, with the exception of restriction cloning scars and few residues at the C-terminal boundary of the construct (Supplementary Figure 1). Overall, the molecular structure of Vg-CTCK is partially consistent with previously characterized CTCK domains [15], defined by two antiparallel beta sheets, each constituted by beta strands arranged in an antiparallel fashion and tightly interconnected through three disulfide bonds located at the boundaries of the beta sheets (Figure 2A-B). A unique feature of Vg-CTCK crystal structure is the presence of a 12 amino acids alpha helix located between the two beta sheets in a region typically characterized by high flexibility and lack of secondary structure elements (Figure 2A-B). Comparison with the AlphaFold3-predicted computational model revealed only minor deviations within the N-terminal region and the C-terminal region, whereas the overall β-strand assembly and the α-helix positioning were fully consistent (Supplementary Figure 2A). All cysteine residues within the domain were found to participate in intramolecular disulfide bonds forming the pairs Cys2000-Cys2029, Cys2016-Cys2043, Cys2025-Cys2083 (Figure 2C). Consistent with data collected in solution, the crystal packing observed in Vg-CTCK experimental structure supports only monomeric arrangements for this protein domain (Supplementary Figure 3).

**Figure 2.**
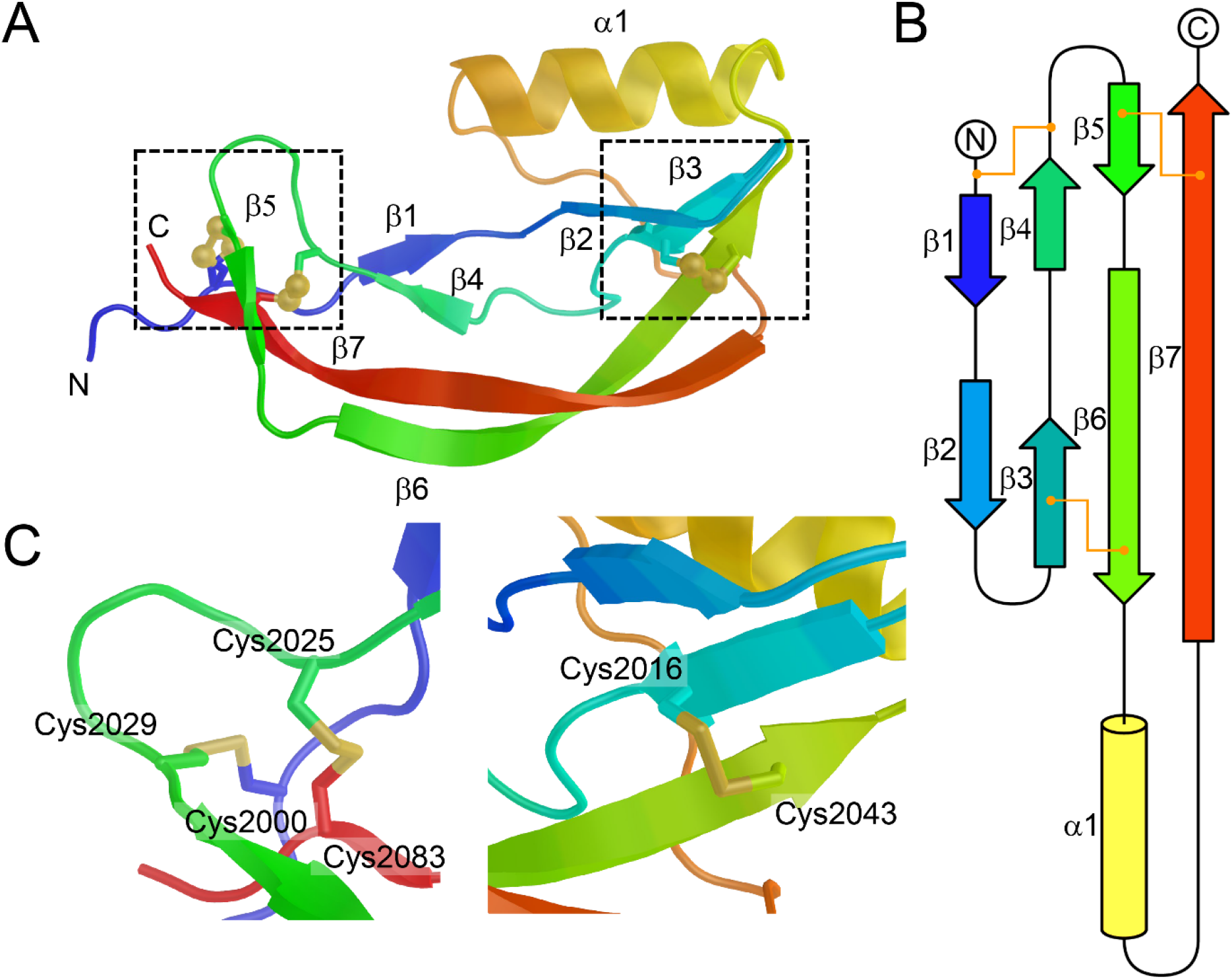
Molecular structure of Vg-CTCK. (A) Cartoon representation of Vg-CTCK molecular structure colored from blue (N-terminus) to red (C-terminus). Secondary structure elements are shown with labels. Disulfide bonds are shown with ball-and stick highlight. The dashed boxes represent the regions shown in the zoomed insets in panel (C). (B) Topology diagram of Vg-CTCK, using the same coloring as in panel (A). Disulfide linkages are shown with orange lines. (C) details of disulfide bonding network (shown as sticks) found in Vg-CTCK. The two panels include the regions highlighted by dashed boxes in panel (A). For clarity, the alternative conformation of the disulfide linkage involving Cys2000 and Cys2029 is not shown.

### Ex vivo crosslinking experiments support absence of Vg-CTCK dimerization

To further explore the molecular features identified in Vg-CTCK structure, we decided to characterize this domain in its native context using cross-linking mass spectrometry (XL-MS). For this purpose, mosquito ovaries were dissected 24 h after the blood meal and immediately cross-linked in phosphate buffer at pH 7.4 containing disuccinimidyl dibutyric urea (DSBU) cross-linker. DSBU is an electrophilic cross-linker that primarily targets lysine side chains and protein N-termini through its NHS ester reactive groups. The samples were then subject to experiments. In our XL-MS experiments, all three *Aedes albopictus* vitellogenins were identified, all of them were found containing the Vg-CTCK domain (Supplementary figure 4). Importantly, the Vg-CTCK domain was detected with 93% to 98% sequence coverage in all three vitellogenins (Table 2), and the three Vg-CTCK domains shared the same amino acid length. Within this domain, vitellogenins display a very high degree of sequence conservation (Supplementary Figure 2B). Since the crystal structure was obtained for the recombinant C-terminal domain of vitellogenin according to sequence XP_019542945.3, we assumed that the corresponding domains in the other vitellogenins would adopt the same overall fold, as suggested by computational predictions using Alphafold3 [21] (supplementary figure 2C). The structural similarity among the different isoforms is further supported by identical cross-links observed in different CTCK domain isoforms, despite slight sequence variations. Specifically, the same residue pairs were found to be cross-linked in two cases: once in XP_019542943.2 and XP_019542959.3, and once in XP_019542959.3 and XP_019542945.2. In both instances, the cross-links mapped to equivalent sequence positions, reinforcing the conservation of the overall fold across different vitellogenins. Accordingly, cross-links identified in homologous C-terminal domains were interpreted in the context of the available crystal structure by mapping the cross-linked residues to equivalent sequence positions and measuring the corresponding Cα–Cα distances.

**Table 2.** Details of *Aedes albopictus* vitellogenin CTCK domain sequences.

|  |  |  |  |
| --- | --- | --- | --- |
| **NCBI entry** | XP_019542945.3 | XP_019542959.3 | XP_019542943.2 |
| **UniProt entry** | A0ABM1Z0L2 | A0ABM1Z1K2 | A0ABM1YVL9 |
| CTCK Sequence start | 1998 | 2069 | 2105 |
| CTCK Sequence end | 2087 | 2159 | 2195 |
| CTCK Sequence length | 89 | 90 | 90 |

Despite the complexity of the *ex vivo* sample, we identified 8 intra-protein cross-links within the C-terminal domain of vitellogenin, and no inter-protein cross-links suggestive of dimerization (Figure 3A). Overall, when mapped onto the Vg-CTCK crystal structure, the cross-link-derived restraints showed strong agreement with a free Vg-CTCK monomer. The Cα–Cα distance distribution of cross-linked lysine pairs was distinct from that of randomly selected lysine pairs from the same structure (Figure 3B). In addition, all cross-links fell within the expected 30 Å threshold of DSBU. Among these, two of the identified cross-links originated from Vg-CTCK isoforms different from the crystallized Vg-CTCK fragment. When superimposed to the experimental Vg-CTCK structure, these crosslinks yielded consistent structural outcomes (Figure 3C), supporting conservation of the CTCK fold across vitellogenins as well as the presence of a folded alpha-helical segment unique of Vg-CTCK. Taken together, XL-MS results of intact ovaries supports the conclusion that the recombinant Vg-CTCK structure is representative of the native fold of this vitellogenin domain and that this domain does not appear to be involved in inter-molecular interactions.

**Figure 3.**
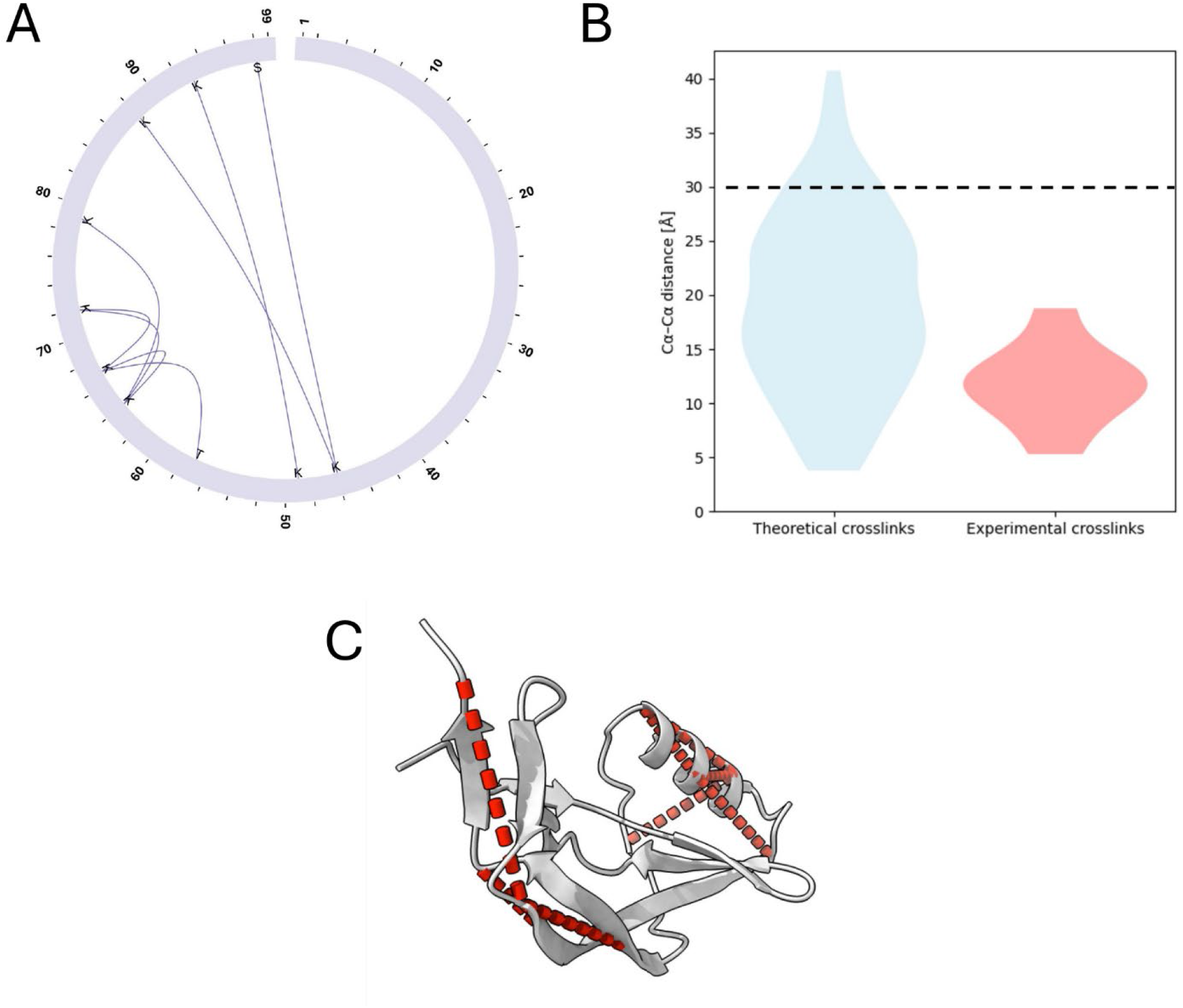
Cross-linking mass spectrometry analysis of Vg-CTCK from *ex vivo* samples. (A) Circos plot showing protein cross-links identified within the C-terminal domain of vitellogenin. (B) Distribution of Cα–Cα distances for experimentally observed cross-links compared to the theoretical distribution of all possible cross-links. The dashed line indicates the 30 Å distance threshold compatible with DSBU cross-links. (C) Cross-links mapped onto the X-ray crystal structure of the recombinant C-terminal domain. Cross-links identified in homologous vitellogenins were mapped onto equivalent residue positions.

### Comparison of Vg-CTCK with homologous CTCK protein structures

Comparative analyses carried out using structural homology softwares yielded very limited similarities with other CTCK protein structures, mostly for the overlapping beta sheets interlocked by disulfide bonds. Using *DALI* [32], the closest match identified was with the structure of the vascular endothelial growth factor (VEGF, PDB ID 2XV7), but the Z-score was extremely low and the superposition RMSD of 4.1 Å suggested very limited similarity. Notably, no structural matches could be identified for the regions involved in VEGF dimerization (Figure 4A). Interestingly, when sorting the *DALI* results based on superposition RMSD, the NMR structure of the human chemokine domain of Fractalkine stood out (PDB ID 1B2T), due to the striking match of a short portion of the beta sheet segment shaped by beta strands β2, β3 and β6, and the conformation adopted by the alpha helix in both proteins (Figure 4B). Notably, the chemokine domain of Fractalkine does not belong to the CTCK domain family. Using *Foldseek* [33], the only reliable (albeit distant) structural matches (probability threshold 0.40) included the CTCK dimerization domain of human Von Willebrand factor (PDB ID 4NT5) and human sclerostin (PDB ID 6L6R, 2K8P). Regarding the Von Willebrand factor CTCK domain, it should be noted that its similarity with Vg-CTCK is limited to the core beta sheet portion, and that the segment involved in disulfide-mediated dimerization in the Von Willebrand factor CTCK domain is not present in Vg-CTCK (Figure 4C). Concerning Sclerostin, similar to what we observed for VEGF, the structural similarity is limited to the core beta sheet portion. Interestingly, Sclerostin is known to act as a ligand of receptors of low-density lipoprotein receptor-related family, such as LRP5/6 [34,35], which share domain architectures with vitellogenin receptors [36,37], however the segment involved in sclerostin-LRP5/6 binding has no matching sequence nor three-dimensional structure in Vg-CTCK (Figure 4D).

**Figure 4.**
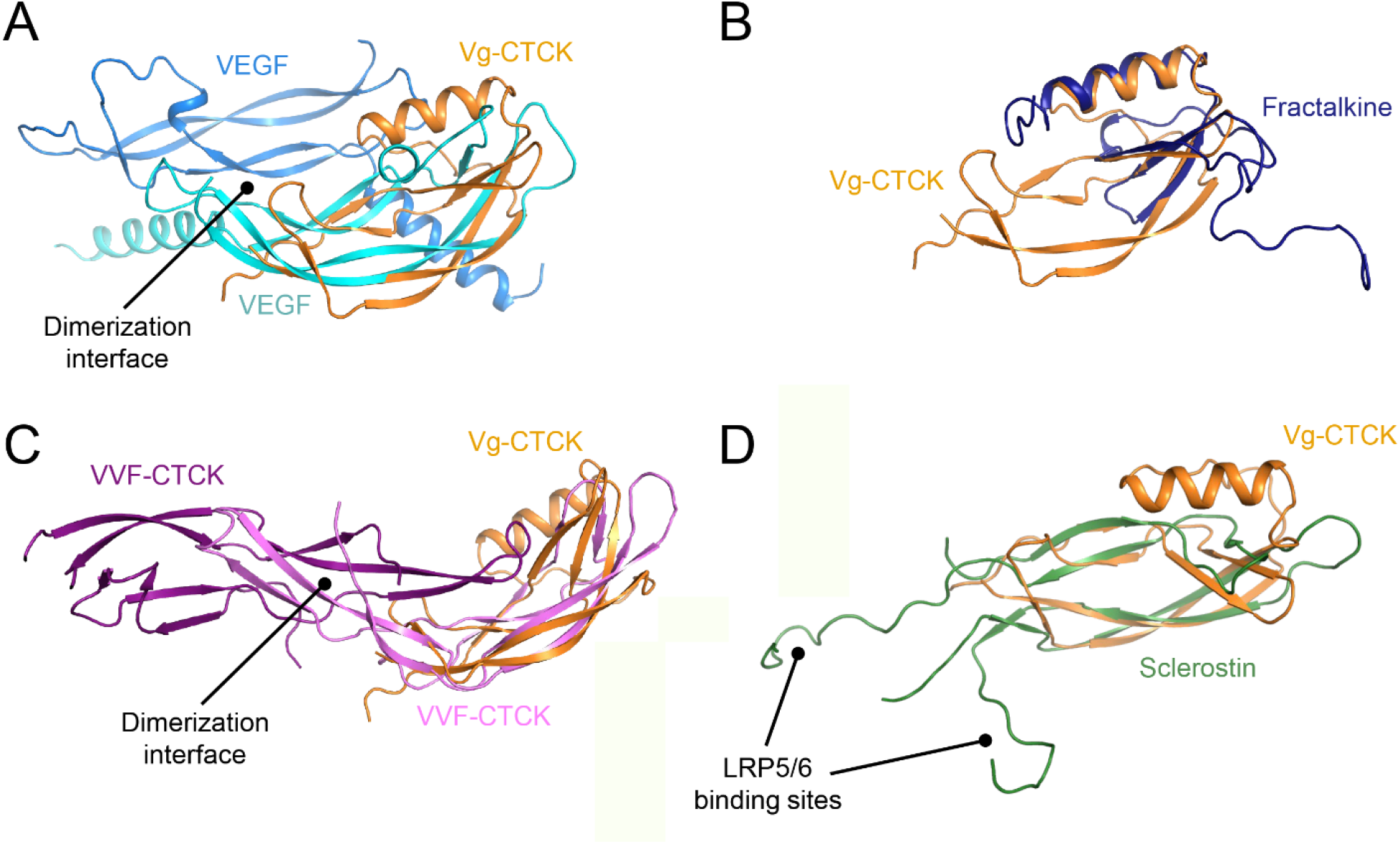
Comparison of Vg-CTCK with other CTCK structures. The panels show superpositions of Vg-CTCK (shown as orange cartoon) with structural homologs identified by *DALI* and *Foldseek*, including dimeric VEGF (A), the chemokine domain of Fractalkine (B), dimeric VWF-CTCK (C), and sclerostin (D).

## Discussion

The vitellogenin CTCK domain is among the most obscure portions of this entire glycolipoproteins. This region displays relatively low tolerance to naturally occurring SNPs in honeybees [38], suggesting possible functional implications underpinned by evolutionary conservation. Several hypotheses regarding the Vg-CTCK have been proposed based on *Apis melifera carnica* data, obtained through nanopore sequencing and subsequent mapping of non-synonymous single nucleotide polymorphism (SNPs) [38], as well as through structural predictions using computational tools [12,16,21]. In honeybees only two common Single Nucleotide Polymorphisms (SNP) are located within the CTCK domain (G1678S and T1676S), and do not pose a drastic change in amino acid sequence, while another common SNP (L1670S) is located within the linker region connecting the C-Terminal domain to the remainder of vitellogenin molecule [38]. Conversely, the C-terminal domain could not be detected in purified honeybee haemolymph preparations, leading to extensive speculations about its role and involvement in vitellogenin maturation through proteolysis [12], mostly guided by the distant structural homology with VWF-CTCK, which is critical for protein multimerization [14]. Based on these observations, it has been hypothesized that the vitellogenin C-terminal domain may participate in dimerization processes and the formation of physiological concatenamers. However, these models currently lack experimental validation [12]. Alternatively, it was proposed an immune-related functions in insects [12]. Despite these hypotheses, the molecular role of the vitellogenin C-terminal domain remains unclear. Although predictive models represent a powerful tool for structural investigation, the limited availability of experimentally resolved structural homologs may reduce the reliability of computational predictions for poorly characterized protein regions.

In this work, we have recombinantly produced the CTCK domain of a vitellogenin isoform of *Aedes albopictus*, performed structural characterization using X-ray crystallography, and combined the structural insights with cross-linking mass spectrometry (XL-MS) investigations on *ex vivo* mosquito samples. The structural data revealed the expected CTCK molecular architecture, defined by elongated beta strands interlinked through an intricate network of disulfide bonds. An unusual alpha helix between strand β6 and β7 constitutes a distinguishing element of Vg-CTCK, unprecedented in experimentally determined structures of this domain family. Analysis of the crystal packing, as well as biophysical characterizations carried out in solution using SEC-MALS ruled out the possibility of homo-dimerization, suggesting that Vg-CTCK is unlikely to contribute to vitellogenin multimerization, as previously proposed [12]. Structural homology investigations further highlight the unique architecture of the Vg-CTCK molecular structure, suggesting distant similarity with extracellular growth factors typically involved in signal transduction cascades rather than dimerization. In this respect, the similarity identified with human sclerostin is particularly relevant, as sclerostin binds receptors that share structural homology with vitellogenin receptors [34,35]. Nevertheless, superpositions of Vg-CTCK and sclerostin highlight that the region involved in receptor binding is not present in Vg-CTCK. This is consistent with our finding regarding the similarity with the CTCK of human VWF, structurally matching the beta sheet arrangement of Vg-CTCK with the exception of the dimerization site featured in VWF-CTCK [14].

Cross-linking mass spectrometry (XL-MS) provides an orthogonal way to test whether a structural model is compatible with residue proximities observed in the native biological context. Because cross-linkers impose defined distance restraints, cross-linked lysine pairs are expected to fall within an upper distance threshold, typically below ∼30 Å when spacer length, side-chain flexibility, and structural uncertainty are considered [39]. We therefore applied XL-MS directly to intact mosquito ovaries to assess whether the X-ray structure of the recombinant C-terminal domain of vitellogenin reflects its conformation in the native organ environment. The absence of inter-protein cross-links between two vitellogenin molecules further argues against stable homodimer formation under these conditions, suggesting that the C-terminal domain is monomeric in the ovarian environment.

In conclusion, our structural, biophysical and *ex vivo* data consistently highlight a monomeric assembly for Vg-CTCK, and do not support the hypothesis of stable, non-covalent dimerization under the tested conditions. Nevertheless, additional experimental investigations will be needed to address whether the Vg-CTCK domain contributes to higher-order vitellogenin assembly formation *in vivo*, or if instead it contributes to other functions in insects triggering unprecedented signal transduction cascades.

## Acknowledgements

We thank the European Synchrotron Radiation Facility (ESRF) for the provision of synchrotron radiation facilities within the macromolecular crystallography beamtime allocation group (BAG) MX-2691 and MX-2790.

## Data availability

Coordinates and structure factors have been deposited in the Protein Data Bank under accession code 31CC. Raw diffraction data are available through the ESRF data portal under accession code https://doi.esrf.fr/10.15151/ESRF-ES-2019878517. All data needed to evaluate the conclusions in the paper are present in the paper or in the Supplementary Materials.

## Funding

This research was funded by the Italian Ministry of University and Research (MUR) and the European Union Next Generation EU (PRIN 2022 PNRR—Project P20224WAME to CI, PG, FF).

## Author Contributions

CI, PG and FF jointly conceived the project and designed research. MB and MS produced and crystallized recombinant Vg-CTCK. MB and FF and performed X-ray data collection, solved the structure, carried out structural refinements, and analyzed the data. PG and MV prepared *ex vivo* mosquito samples. AV, AA, FM, and CI produced and analyzed the proteomics and XL-MS data. MB, FF, PG, and CI wrote the paper, with contributions from all authors.

## Conflict of Interest statement

The authors declare no competing financial interests.

## Notes

### Competing Interest Statement

The authors have declared no competing interest.

