## Supplementary Information for "Structural studies of the C-terminal cysteine-rich domain of *Aedes albopictus* vitellogenin reveal an evolutionarily divergent, monomeric C-terminal cysteine knot (CTCK) architecture"

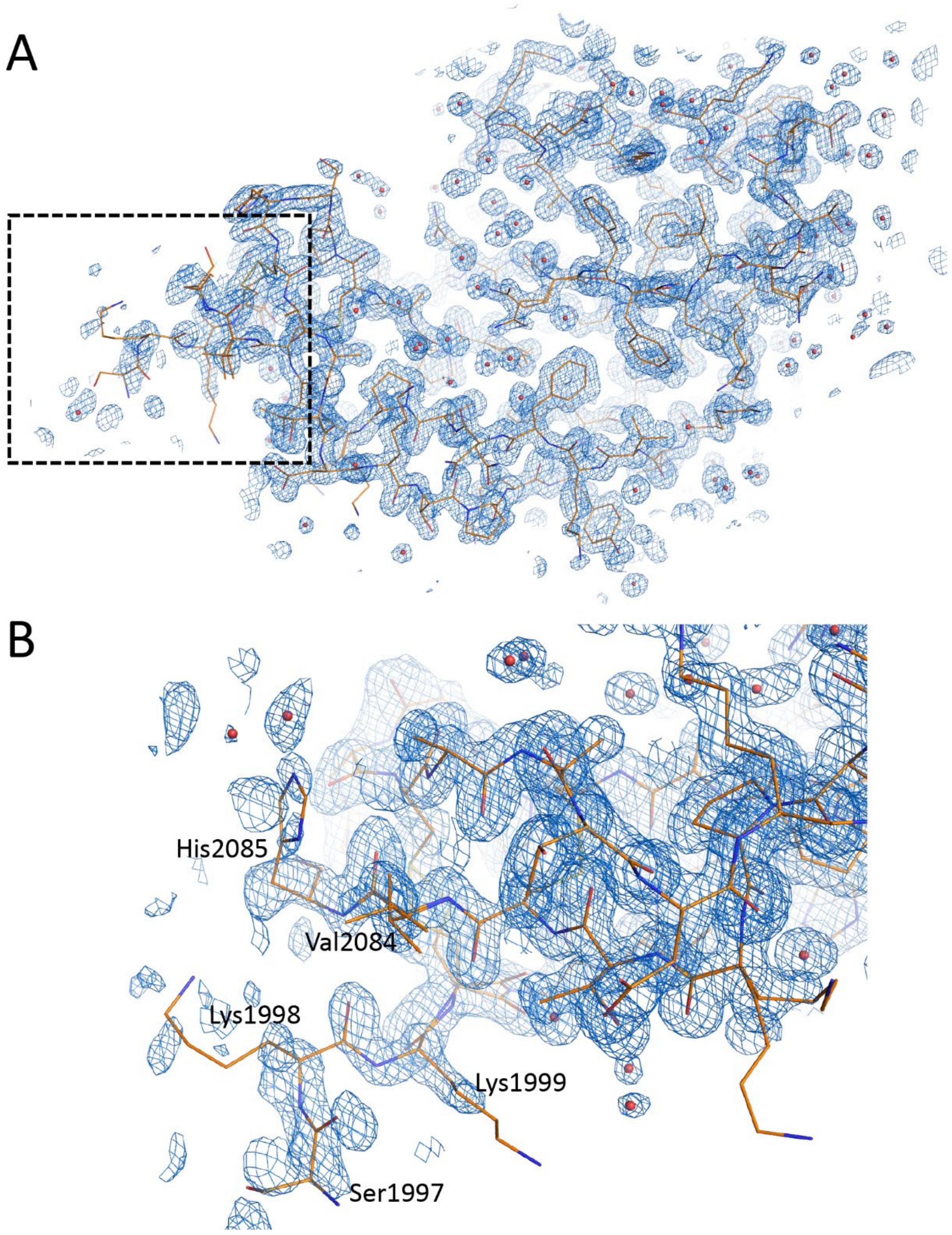


**Supplementary Figure 1. Quality of the experimental electron density in the Vg-CTCK crystal structure.** (A) Refined $2F_o$-$F_c$ electron density map (blue mesh, contour level 1.2 $\sigma$) covering the entire Vg-CTCK domain (shown as orange sticks). Water molecules are shown as red spheres. The dashed box shows the location of the N- and C- terminal segments, highlighted in the detail shown in (B). Modelled residues located at the N- and C- boundaries are labeled.

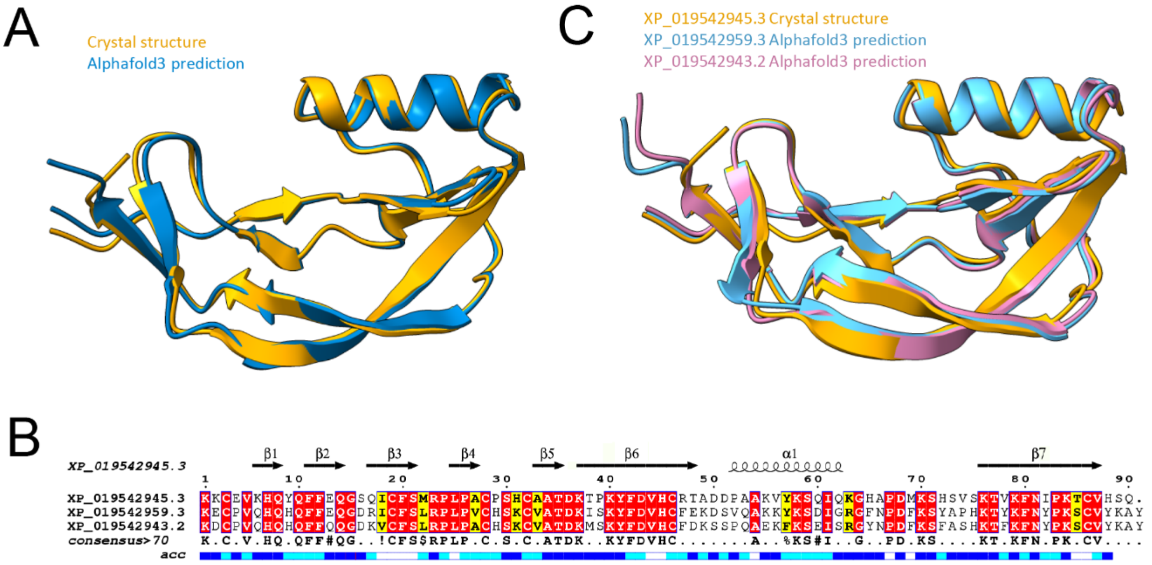


**Supplementary Figure 2. Comparison of computational models and crystal structures of Vg-CTCK.** (A) Superposition of experimental (orange) versus theoretical (blue) structures of *Aedes albopictus* Vg-CTCK. (B) Sequence alignment of Vg-CTCK domains from the different *Aedes albopictus* vitellogenin isoforms. The “consensus” line, derived from the alignment software, indicates conservation of sequence identity or similarity. The “acc” line indicates solvent accessibility (white = not accessible, blue = solvent accessible). (C) Superposition of predicted Alphafold3 models of Vg-CTCK from different *Aedes albopictus* vitellogenin isoforms s with the experimental structure of Vg-CTCK.

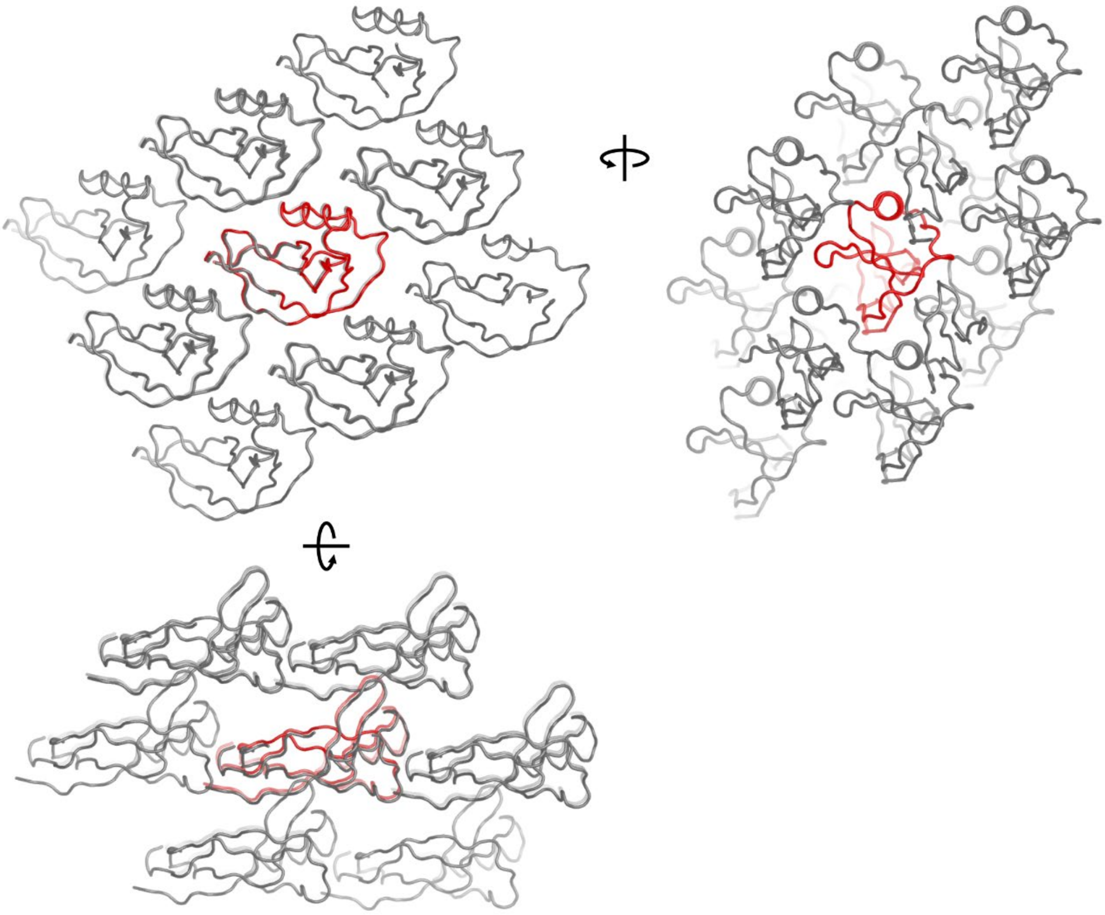

**Supplementary Figure 3. Overview of the crystal packing network observed in Vg-CTCK structure.** The molecular structures of Vg-CTCK are shown with ribbon style. To facilitate identification of the repeating element, the Vg-CTCK constituting the asymmetric unit is colored in red, whereas neigboring molecules are colored in grey. The representation is provided in three different orientations, based on the triclinic unit cell.
